# Polycomb repression of Hox genes involves spatial feedback but not domain compaction or demixing

**DOI:** 10.1101/2022.10.14.512199

**Authors:** Sedona Murphy, Alistair Nicol Boettiger

## Abstract

Polycomb group (PcG) proteins modulate higher-order genome folding and play a critical role in silencing transcription during development. It is commonly proposed that PcG dependent changes in genome folding, which compact chromatin, contribute directly to repression by blocking binding of activating complexes and demixing repressed targets from non-repressed chromatin. To test this model we utilized Optical Reconstruction of Chromatin Architecture (ORCA) to trace the 3-dimensional folding of the *Hoxa* gene cluster, a canonical Polycomb target, allowing us to analyze thousands of DNA traces in single cells. In cell types ranging from embryonic stem cells to brain tissue, we find that PcG-bound chromatin frequently explores decompact states and partial mixing with neighboring chromatin, while remaining uniformly repressed, challenging the repression-by-compaction model. Using polymer physics simulations, we show that the flexible ensembles we observe can be explained by dynamic contacts mediated by multivalent interactions that are too weak to induce phase separation. Instead, these transient contacts contribute to accurate propagation of the epigenetic state without ectopic spreading or gradual erosion. We propose that the distinctive 3D organization of Polycomb chromatin, reflects a mechanism of “spatial feedback” required for stable repression.

## Introduction

Polycomb group (PcG) proteins form multi-subunit complexes that interact with chromatin to repress transcription. This repression needs to be both robust and reversible, particularly during development where correct spatiotemporal gene expression patterns are controlled by balanced activating and repressive signals. Biochemical interactions between the Polycomb (Pc) complexes and the chromatin regions they target, reinforce maintenance of repression-associated chromatin states throughout development (epigenetic memory) while also altering their 3-dimensional genome organization (*1–6*). This reorganization of the 3D genome is widely described as a compaction event that underlies repression of Pc-targets (*1–6*). Recent experiments and models have proposed that liquid-liquid phase separation (LLPS) of PcG proteins may underlie this compaction and further segregate Pc-domains from neighboring active chromatin (*7–10*).

Compaction of Pc-bound DNA has been measured across chromatin length scales. First observed in *in vitro* nucleosome compaction assays (*11*), the compaction phenotype engendered a model for Polycomb-mediated repression in which the formation of compacted domains excludes transcription machinery, thus creating a structural barrier to activation. Since then, a variety of sequencing and imaging based techniques have been applied to measure this compaction phenotype on length-scales from 1s-100s of kilobases and largely support the structural barrier model. DNA-FISH based assays have shown that when *Hox*-gene complexes lose Polycomb repression upon activation, the average distance between the ends of the domain increases (*12–14*). Superresolution microscopy techniques have demonstrated that the average Polycomb domain occupies a smaller volume than other non-Pc bound, silent chromatin or active regions (*15–19*). Chromatin conformation capture assays that use a sequencing readout, like Hi-C, have shown that Polycomb-bound regions form compact domains that decompact as a domain activates (*18, 20, 21*).

From these studies it is clear that Polycomb-bound regions adopt unique structural features that appear to be important for gene repression. However, it is not well understood how variable domain folding is within compacted regions across cell populations or time. Technical challenges limited previous analyses of variability in compaction state and domain folding. Bulk sequencing assays (Hi-C) lack the cellular resolution to assess variability, while single-cell sequencing approaches lack the sensitivity to assay structures at the scale of most Polycomb-domains (a few to few hundred kb). Meanwhile, superresolution microscopy techniques like STORM/PALM, are limited in capturing the polymeric nature of chromatin because they provide only volumetric or density measurements. In recognition of these uncertainties, prior super-resolution microscopy work, including our own, focused on population-level properties such as average domain size and average separation, even though the image data is intrinsically single-cell.

To address these limitations, we utilized Optical Reconstruction of Chromatin Architecture (ORCA) ((*22–24*)), to trace the 3D structure of the *Hoxa* gene cluster, a canonical Polycomb target, in thousands of single-cells. By quantifying the relative compaction and demixing of single-cell DNA traces we find that the DNA folding mediated by PcG proteins does not stably compact or de-mix chromatin. While the *Hoxa* locus is statistically more likely than a neighboring region to adopt a more compact and de-mixed structure, it is flexible and varied, challenging the structural barrier hypotheses of compaction and phase-separation. By performing molecular-dynamics polymer simulations, we construct an alternative model linking the observed flexible 3D structures of Polycomb-chromatin to repression.

## Results

While Polycomb-bound regions have been extensively studied via super-resolution microscopy by us and others (*15–18*), several technical limitations have made understanding the variability of folding, compaction, and separation at single-cell resolution difficult. First, single or two color super-resolution images provide a limited view of how the sequence is folded or how uniform or variable the compaction is across the domain, (**Fig. 1A)**. Variation in label efficiency (at the single-molecule level) contributes to variation in probe density across cells - a technical variability not readily distinguished from biological variation in chromatin density. Intrinsic single-molecule variation in the number of photo-cycles per fluorophore and the photon count per emission cycle further compound this problem – a single fluorophore may bleach before blinking or may blink 100 times before bleaching (*25*), so localization density is not a perfect measure of probe density. Arbitrary thresholds are needed to distinguish background signals from on-target signals. These thresholds are more likely to mistake low-density regions as background and may overestimate compaction **(Fig. 1A)**. A systematic over-estimation of compaction of Polycomb-bound regions would reinforce a model of structural inaccessibility mediated by domain folding.

**Fig. 1.**
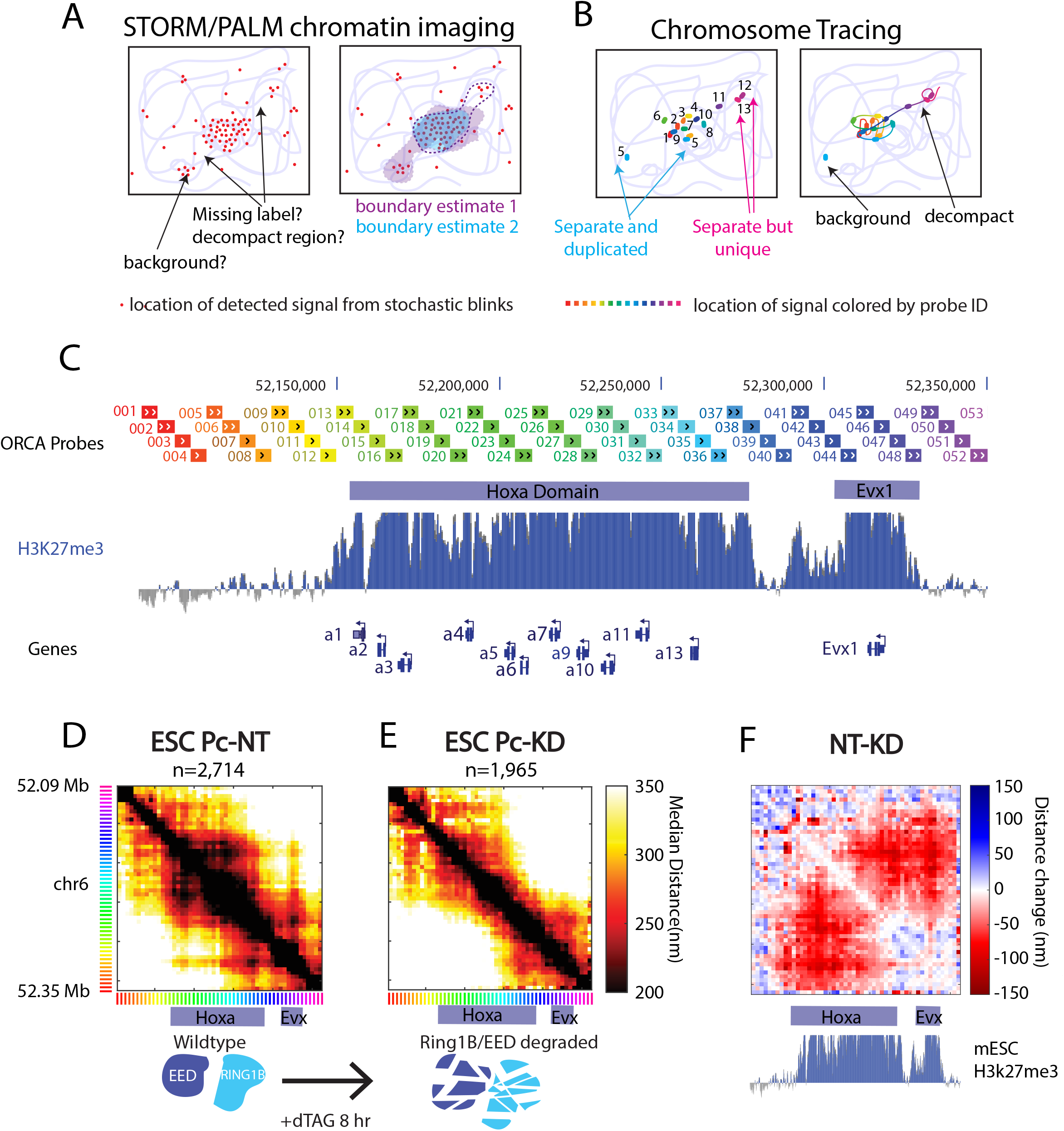
Advances in imaging technology improve measurements of 3D chromatin structure and quantification of Polycomb-dependent compaction. **(A)** Schematic of the uncertainties of common single-molecule localization microscopy (SMLM) super-resolution techniques, such as STORM/PALM. Red points mark fluorophore locations identified during stochastic blinking. Blue lines denote chromatin. **(B)** Schematic of the advantages of chromosome-tracing imaging techniques. Rainbow dots mark locations of sequentially imaged genomic segments, as shown in a genome browser in C. **(C)** Genome browser track annotating the Polycomb domains, H3K27me3 repressive mark, 5 kb ORCA probes, and genes. **(D)** Median distance map of Hoxa locus from untreated mES cells carrying RING1B and EED degrons. **(E)** Median distance map from mES RING1B+EED degraded cells after 8 hr of degron induction with dTAG ligand. **(F)** Difference of the maps shown in (D) and (E).

The advent of sequential imaging technologies for chromosome tracing (*26*) addressed the limitations discussed above, and recent improvements to achieve kilobase resolution now enable its application to finer-scale chromatin features like Polycomb domains (*19, 27, 28*). These methods use sequential addition and removal of fluorescent probes to trace the chromatin path in chemically fixed cells. Probes from each step in the path are identified by unique labels, and thus do not require arbitrary density thresholds to determine if they are part of the domain or not. For example, if probe 5 gives an extraneous foci outside the domain, in addition to on-target foci inside the domain, the true probe 5 may be distinguished from the background probe 5 by its distance to other probes (**Fig. 1B**). If probe 12 gives a foci outside the domain because the intervening sequence is decompact, this case may be distinguished from the background event just considered by the absence of probe 12 signal around the other probes (**Fig 1B**). The known copy number of the locus (which is also measured by the hybridization) provides valuable information that removes the need for arbitrary density or brightness thresholds which may bias estimates of compaction. The labeling efficiency is measured per cell, per step along the domain, rather than being an unknown quantity (**Fig. 1B**). In addition to the improved level of structural detail provided by a trace rather than a volumetric region (blob), these additional features encouraged us to investigate the single-cell variability in structure of Polycomb-repressed chromatin with ORCA.

We focused our studies on the *Hoxa* cluster, a ∼100 kb region containing homeobox genes with critical roles in anterior-posterior patterning, that were the first identified targets of the Polycomb system and one of the best-studied examples of Polycomb-associated repression and memory (*29*). This domain is demarcated by the H3K27me3 repressive mark, which is deposited by PcG proteins and is required for repression (*30*). Primary probes were designed to span the *Hoxa* locus at 5 kilobase resolution, extending upstream and downstream of the cluster, to total 260 kb of coverage **(Fig 1C)**. Imaging of the *Hoxa* cluster produced single-cell snapshots where each detected region can be connected in linear genomic order to reconstruct a 3-dimensional (3D) trace. Population pairwise-contact frequency, computed from these single-cell traces, correlated well (*r* = 0.95) with deep in-situ Hi-C data from recent studies (*31*) (**Supp Fig. 1**). To determine which features of the 3D structure are dependent on Polycomb-activity, we utilize the dTAG degron system (*32*) to acutely deplete RING1B and EED (*33*), core, catalytic subunits of the Polycomb system in mouse embryonic stem (mES) cells. The depletion of these two PcG proteins is sufficient to abolish the Polycomb repression (*33*). This degradation is fast and no protein is detectable by immunofluorescence or Western blot after 8 hours of dTAG treatment (**Supp Fig. 2**). This allows for direct measurement of Polycomb-specific features that are not confounded by secondary effects of long-term Polycomb withdrawal, in contrast to earlier approaches using siRNA (*15, 34*) or gene deletions (*12, 13, 18, 20, 21, 35*).

Deactivation of Polycomb repression with dTAG induced substantial changes in the average 3D organization of the *Hoxa* cluster, seen in the comparison of the median pairwise distances throughout the domain (**Fig. 1D-F**). Median pairwise distance maps were reproducible between experiments, and showed similar changes in 3D structure between replicates (**Supp. Fig. 3A,B**). The domain borders upstream and downstream of the complex, which aligned to the H3K27me3 region in wildtype cells, became less pronounced following degradation (**Fig. 1D,E**). The border downstream of *Hoxa13* was replaced with a new border near the middle of the *Hoxa* complex (**Fig. 1C-E**). The average separation between *Hoxa* and *Evx1* increased, removing the stripe of contacts which in wildtype cells extends from *Evx1* through *Hoxa1* (**Fig. 1D,E**). As the distances in these maps are measured in nanometers (not relative reads), and the induced and uninduced cells were imaged together in the same experiment to avoid batch effects, these average distance maps can be subtracted from one another (**Fig. 1F**). This reveals a significant expansion throughout the H3K27me3 marked region (see **Supp. Fig. 3C** for p-value map) with much of the domain an average of over 100 nm further apart (**Fig. 1F**). This significant increase in the average separation throughout the domain aligns well with previous observations (*12–14, 16, 17, 19, 21, 36*).

While aggregate data is consistent with the previously proposed models of compaction and/or phase separation of the *Hoxa* domain as a physical mechanism for repression, we examined individual 3D traces to determine if they adopted physically compact and separate folding patterns relative to neighboring chromatin (**Fig. 2A**). We found that individual loci revealed a diverse array of structures, many of which appeared neither compact nor well segregated from neighboring chromatin (**Fig. 2B-I**). To quantify the degree of compaction population wide, we calculate the radii of gyration (R_g_), which is the distance of all measured points in the trace from the center of mass of that trace, and an ideal metric of polymer compaction (*37*). We compared R_g_ of the Polycomb-repressed *Hoxa* region and the Polycomb-free region immediately upstream of *Hoxa* **(Fig. 2A**). The upstream region acts as an internal control for cell cycle, cell size, and other potentially confounding factors. R_g_ distributions across cell populations were reproducible across replicate experiments (**Supp. Fig 4A,B**). Comparing the R_g_ in treated vs untreated cells, we observe substantial variation in both Pc-domain and non-Pc domain R_g_ across the population (**Fig. 2C,D**). While the R_g_ ratio (compaction ratio) of non-Pc to Pc chromatin is on average greater than 1 in untreated cells (as expected for a compact domain) and shifts close to 1 in treated cells, the distributions overlap substantially (**Fig. 2E, Supp. Fig 4E**, note the figure shows log-ratios). The degree of compaction in a given cell at the instant of fixation is therefore weakly indicative of its Polycomb state. In fact, only 28% of untreated traces had Pc-chromatin more than twice as compact as the neighboring chromatin.

**Fig. 2.**
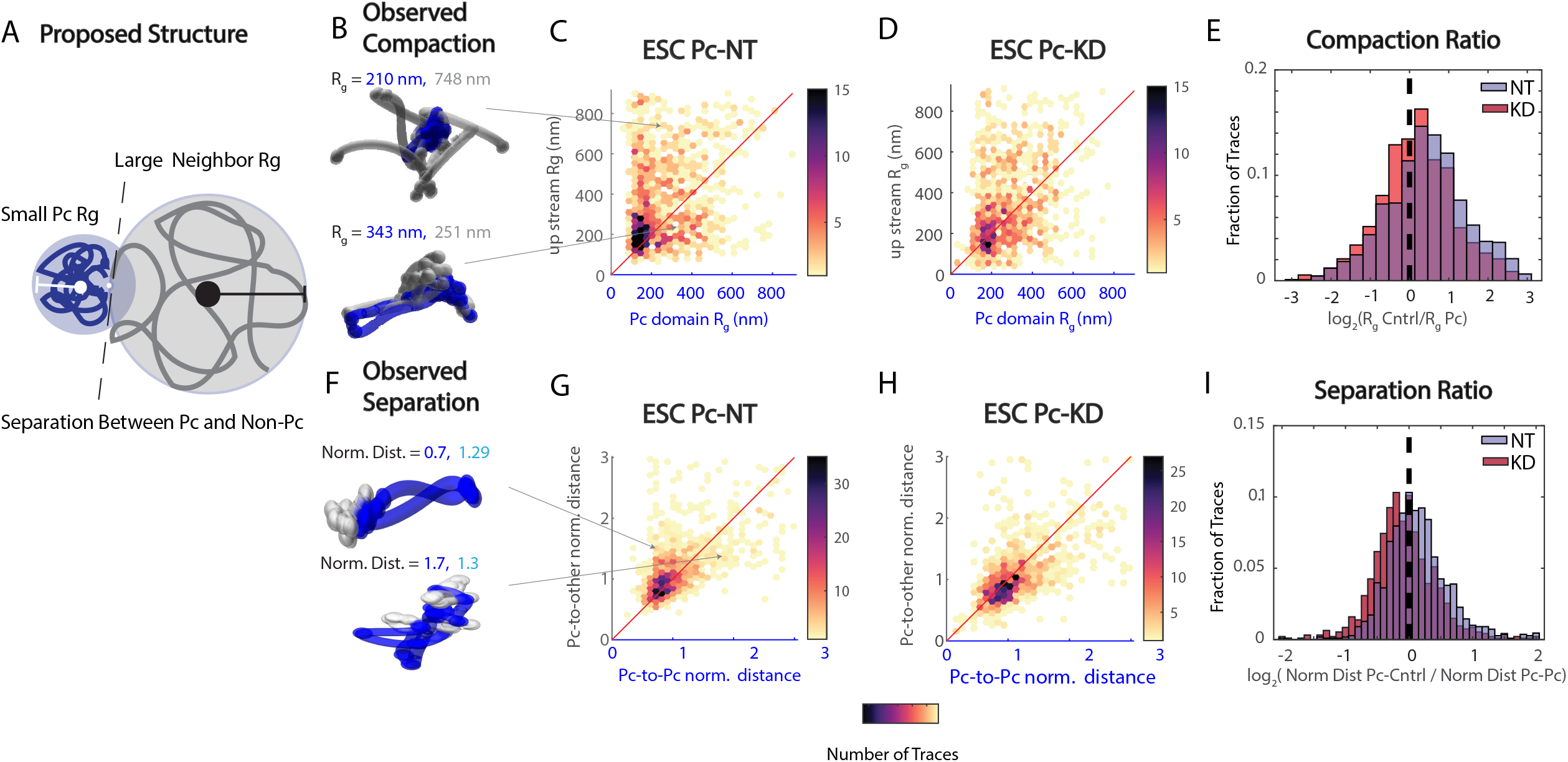
The Hoxa locus in mES cells is frequently de-compact and un-separated. **(A)** Cartoon of the predicted structure for Pc-domains, illustrating the metrics to quantify compaction (relative radius of gyration, Rg), and separation of domain. **(B)** Example traces of the Hoxa region in mES cells. Blue denotes the region targeted by the Pc-system Gray denotes neighboring non-Pc targeted region (R_g_ in nm). Arrows point to where polymers exist in C. The R_g_ for the Pc-targeted and upstream non-Pc region are shown in blue and gray text. (**C)** 2D plot comparing the R_g_ of the Polycomb-domain and the R_g_ of the upstream region for each measured loci in untreated mES cells (n=2741). Points above the red line indicate the Polycomb-domain R_g_ is smaller (nm) than upstream-domain R_g_. **(D)** as in B, but for dTAG treated cells (n=1965). **(E)** Compaction ratio plotted as the log2 ratio of the upstream-domain R_g_ to the Polycomb-domain R_g_ for the no treatment and dTAG knockdown samples. **(F)** Example traces of Hoxa region. Arrows point to where polymers exist in C. **(G)** 2D plot comparing the Pc-to-Pc normalized distances to the Pc-to-other normalized distances in each measure loci in untreated mES cells (n=2741). Points above the red line indicate the Pc-to-Pc normalized distances are smaller than the Pc-to-other. Normalization computed how 3D distance scaled as a function of genomic (1D) separation across the entire domain, and divided observed distances by this expected distance. **(H)** as in G, but for dTAG treated mES cells (n=1965). **(J)** Separation ratio plotted as the log2 ratio of the Polycomb-domain normalized distances and the upstream-domain normalized distance for the no treatment and dTAG knockdown samples.

As it has also been hypothesized that demixing (*15, 17*) and/or phase separation (*7, 8, 38*) may underlie Pc-mediated repression, we next quantified separation of individual traces. This separation metric is a distinct structural feature from the compaction ratio (**Fig. 2 B vs F**), though the two properties are moderately correlated as expected (**Supp Fig. 5**). Individual traces from both untreated (**Fig. 2F**) and PcG-depleted cells (**Supp. Fig. 6**) show a varied organization, wherein some cells separated *Hoxa* from neighboring chromatin and others exhibit an intermixed conformation. To quantify separation, we first computed distances between all pairs. We normalized these distances by their ‘expected’ distance given their genomic separation (see **Methods**). We created a 2D graph, where for each single trace, we plotted the average normalized distance of all Pc-to-Pc pairs versus the average of all Pc-to-other pairs (**Fig. 2G,H**). The graph for all traces from untreated cells shows a broad distribution trending to closer Pc-to-Pc vs. Pc-to-other (**Fig. 2G,H**). Examining the ratio of these normalized distances, we found approximately 60% of untreated traces exhibited separation ratios greater than 1 **(Fig. 2I, Supp. Fig. 4F)**. This fraction dropped to ∼40% of traces upon PcG depletion. Thus, while PcG activity clearly affected the frequency with which PcG-chromatin segregated from non-PcG chromatin, it is far from achieving a uniformly demixed or phase-separated state.

To compare heterogeneity in Pc-domain folding to transcriptional states, we next analyzed tissues from cryosectioned mouse embryos **(Fig. 3A,B**), and quantified both transcription and repression state of the cells alongside the structure of the *Hoxa* cluster. Single molecule RNA FISH (smFISH) experiments in posterior embryonic tissues validated our *Hox* RNA probes, with robust staining in the posterior tissues and lack of signal in the most anterior (**Fig. 3A**). In contrast, we detected no notable signal from the *Hox* probes tested in E10.5 embryonic brain sections, validating the uniformly repressed state of the locus in these cells (**Fig. 3B,C**). Positive control RNAs were robustly detected in nearly all cells, indicating the lack of *Hox* RNA signal was not an artifact of RNA preservation (**Fig. 3B,C**).

**Figure 3:**
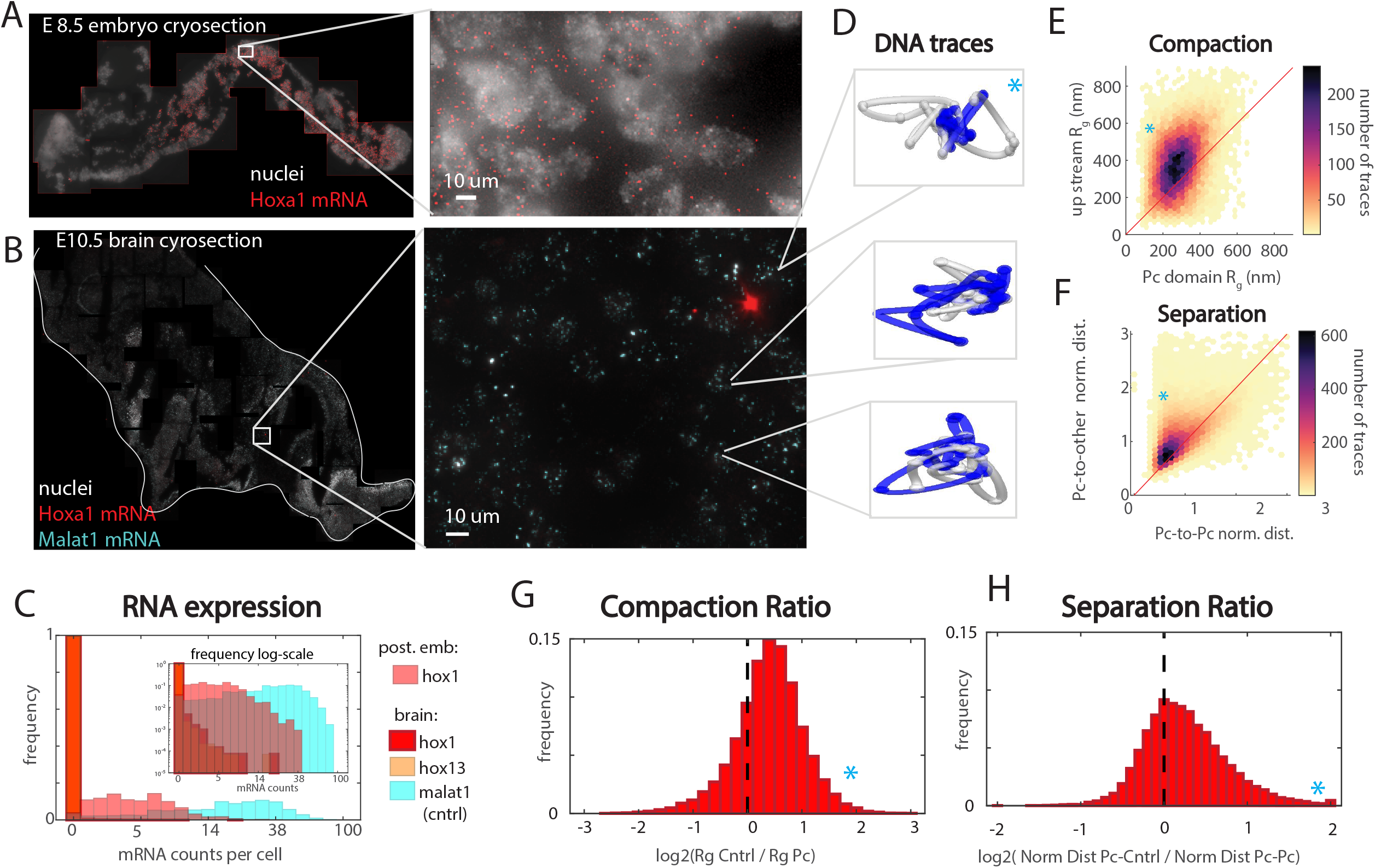
The Hoxa locus in brain tissue is uniformly repressed but not uniformly compacted nor separated. **(A)** E8.5 mouse embryo section labeled with probes targeting *Hoxa1* RNA. **(B)** E10.5 mouse embryo brain section labeled with probes targeting *Hoxa1* and *Malat1* RNA **(C)** Histogram of RNA expression per cell for *Malat1, Hoxa1*, or *Hoxa13* in the E10.5 brain section and Hoxa1 in selected control tissue from the mouse embryo. Wilcoxon rank sum test comparing fraction of hoxa1 silent to fraction compact p=2.5e-34, *Hoxa1* vs fraction separated, p=2.5e-34. **(D)** Example chromatin traces with *Hoxa* domain labeled in blue. **(E)** 2D-compaction plot comparing upstream and Polycomb-domain R_g_ from brain cells, as in Fig. 2B. n= 26,135 traces **(F)** 2D-Separation plot comparing normalized Polycomb-to-Polycomb distances and Polycomb-to-other distances. **(G)** Histogram of the compaction Ratio, as in Fig 2. **(H)** Histogram of the Separation Ratio, as in Fig 2. Blue stars denote where the example trace in D lies relative to the population analyses in E,F,G,H.

We then applied ORCA in cryo-sections from the same two brains used for RNA analysis to look at the 3D structure of the *Hoxa* cluster. As data from both brains showed strong correlation between the datasets, we merged them to increase statistical power (r = .99, **Supp. Fig. 7A,B**). While individual traces exhibiting visually compacted and segregated organization could be readily observed (**Fig. 3D**, blue star), most traces exhibited more open and intermixed organization (**Fig 3D**). By quantifying this as above, we observed the 2D-Rg distribution shift to the upper left compared to mESCs, reflecting smaller Rg for Pc-domains compared to the neighboring region. Nonetheless, many traces (23%) still have Pc-domains that are equal to or larger than neighboring regions (**Fig. 3E, Supp. Fig. 7C,D**). The distribution of normalized distances (separation) also exhibited shorter distances among Pc-to-Pc compared to Pc-to-other (68%), while maintaining a broad distribution across the population (**Fig. 3F, Supp. Fig. 7E,F**). The heterogeneous degree of compaction and separation can be seen more clearly in the ratio of domain R_g_ (compaction ratio) (**Fig. 3G**) and the ratio of the normalized Pc-Pc distances to normalized Pc-to-other distances (separation ratio) (**Fig. 3H)**. This heterogeneous occupancy of the structural states, highly variable in their degree of compaction or separation from neighboring chromatin, contrasted with the uniform repression observed at the transcriptional level (**Fig. 3C**).

We found these observations difficult to reconcile with the current models, in which compaction and/or phase separation directly underlie transcriptional repression. We turned next to a simulation approach, to ask what sort of model could explain the origin of this heterogeneous, apparently flexible 3D folding observed in the data. We then asked if any properties of this alternative model suggested a functional role for flexible domain folding in repression.

We began our modeling by incorporating a minimal number of features of the Pc-repressed chromatin required to produce agreement with our microscopy data (**Fig. 4A-C**). Chromatin was modeled as a floppy chain of beads moving around under the thermal motion in a confined environment (**Fig. 4A**). Beads could be either blue or gray, representing the Pc and non-Pc chromatin states. Blue beads were allowed to stick to one another while gray beads were not. This “stickiness” aimed to capture the observation that multiple PcG proteins have been shown to aggregate in purified isolates, in cells, and/or on chromatin templates (*11, 21, 35, 39, 40*). In development transiently expressed cell type-specific Transcription Factors (TFs) establish Pc-domains and then switch off, relying on the Pc-system to maintain the repression (*5, 29*). To capture this process, color states were assigned at the start of the simulation, but allowed to evolve in time by the following rules. We allowed blue beads to switch to become gray beads, capturing the dynamics of nucleosome turnover and the transient nature of PcG-chromatin binding (*41–43*). Gray beads could also switch to become blue. We assumed this happened spontaneously, at a low frequency, but was catalyzed (i.e. more probable) if gray beads were in proximity to one or more blue beads. This feature was motivated by the known behavior of PcG proteins. Some catalyze post-translational modifications of chromatin (such as H3K27me3) that facilitate binding of other PcG factors, while others recruit each other in H3K27me3 independent means (*1–3, 5, 8*). Meanwhile, the low frequency spontaneous conversion aimed to capture the ability of PcG proteins to interact with naive chromatin at low efficiency (11, 35, 36)(33). Finally, we allowed for 3 different degrees of blue, which must all be lost for the bead to become gray (**Fig. 4B**). These 3 states may be thought of as different methylation states, different sizes of PcG-aggregates, or different states/subunit composition of PcG complex assembly (**Fig. 4C**). Additional states beyond 3 would produce similar results to those described below, but we were unable to capture key features with only a single state. Regulating the increase of state, we assumed the total amount of PcGs in the system to be constant and limiting, so not all beads could be blue at the same time. This was motivated by the sensitivity of cells to PcG dose (*2, 3, 5*), and we found it to be a required assumption to avoid the system becoming trapped in the all-blue state, as recently reported in parallel work simulating chromatin epigenetics (*44*). The 3D polymer dynamics in these simulations were computed using the polychrom framework from open2c (*45*), derived from earlier models of chromatin folding by the Mirny lab (*46, 47*). The recoloring dynamics (modeling PcG biochemical interactions) were added onto this as described in the methods and our deposited code (see **Supplement**), similar to the approach used in recent work (*44, 48, 49*).

**Fig. 4.**
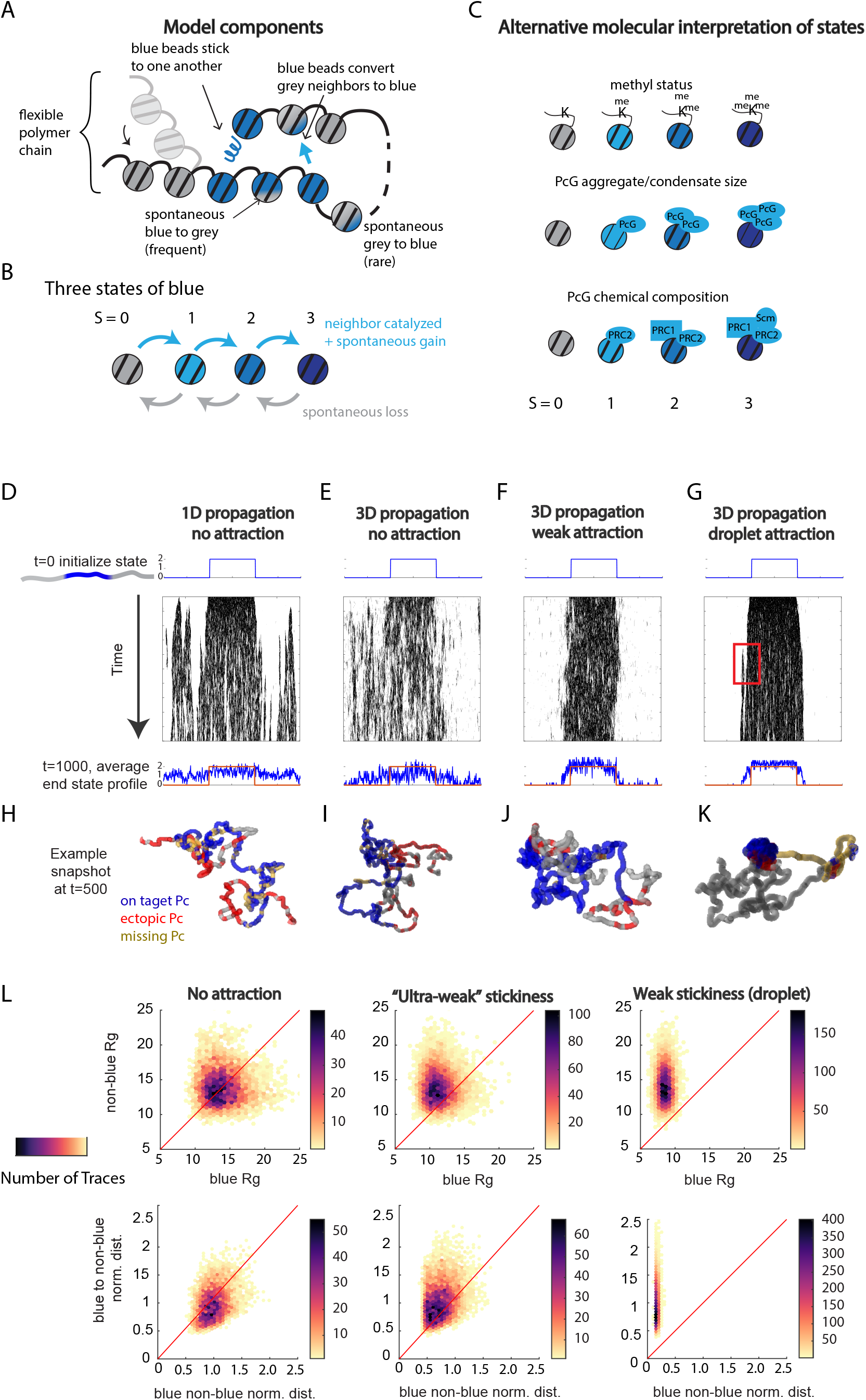
Simulations of PcG spreading and turnover identify how 3D-structure may stabilize repression without forming compact or separated domains. **(A)** Schematic illustrating key parameters of the model (arrows). Blue beads model PcG bound chromatin. (B) Schematic illustrating 4-state model. Gain of state is catalyzed by the presence of neighboring blue molecules and a limiting pool of blue factors (PcG proteins), but does not depend on the beads current state. (C) Schematic depiction of different molecular interpretations of the 4 states shown in B. **(D)** Kymographs from simulations of the 1D propagation model, with no sticky interactions. X-axis reports monomer position (genomic coordinate) and y-axis reports time (arbitrary units). The average starting state and average finishing state across independent simulations are shown above and below the kymograph in blue. The starting state is superimposed in orange on the end state for ease of comparison. **(E)** as in D, but in a simulation allowing state conversions to be catalyzed based on 3D proximity rather than just 1D proximity. **(F)** as in E, but with ultra weak sticky interaction between monomers in states 1-3. **(G)** as in F, but with slightly stronger sticky interaction. Red box highlights a region that converted from PcG free (state 0) to PcG associated and remained so throughout the simulation. **(H)-(K)** Example polymer snapshots at t=500 where blue is on-target PcG, red is ectopic, and yellow is missing PcG, as defined by comparison to the initial state. See also **Supplemental Movies 1-3. (L)** 2D-plots of separation and compaction for the modeled conditions.

We evaluated the stability of repressive domains in these simulations, across a range of conditions, by the ability of the simulation to maintain the originally specified blue-beads in a blue (Pc) state, rather than opening holes of gray beads in the domain or spreading to previously non-blue regions (**Fig. 4D-G**). In this, the model parallels prior work on epigenetic stability in 1D models of the linear genome (*50–54*), though these models intentionally ignore the effects of the 3D domain folding. We also evaluated the simulations on their ability to reproduce 3D organization based on the individual traces (**Fig. 4H-L**) and population average contact frequency (**Supp. Fig. 8**). Simulating the effect of sticky interactions and crowding on 3D structure of Polycomb DNA parallels prior modeling of chromatin folding (*15, 37, 55–59*), though these works assumed an unchanging chemical identity of each bead (changing color/ epigenetic state). The theoretical effects of interactions between changing chemical identities and flexible 3D structure of a polymer has been less explored. The few pioneering works in this area have yet to be applied to data or Polycomb biology (*44, 48, 49, 60*). Notably, we found these simulations that combined spatial and chemical interactions were able to produce behaviors that capture key features of our microscopy data.

We started with the simplest model, in which the gray-to-blue catalyzed addition (PcG spreading) may only happen linearly along the chromatin-polymer backbone. In this model, we find the blue (PcG) domain quickly separated into randomly distributed short islands (**Fig. 4D**). Allowing spreading to occur as long as beads have proximity in 3D, rather than purely 1D, resulted in even more rapid loss of the original pattern (**Fig. 4E**). However, allowing both 3D spreading and a small degree of stickiness among blue beads lead to emergent “epigenetic memory”, with notably more stable propagation of the blue state than the previous models (**Fig. 4F and G**). The sticky interactions make the PcG-bound regions more likely to be in 3D proximity to other PcG-bound regions, and thus more likely to receive replacement PcG when some are lost due to the steady turnover (**Supp. Movies 1-3**).

We observed multiple structurally distinct regimes for different strengths of blue-blue sticky interactions (**Fig. 4H-K**). For all structural calculations of compaction, separation, and pairwise distances, we incorporated an added technical noise to the bead position before the computation. This positional uncertainty added to the simulations was chosen to match the experimentally measured 3D uncertainty from the ORCA traces (see (*19*) and **Methods**). With no stickiness, the 3D organization of the polymers followed the expected behavior for free polymer motion, and blue and gray regions did not exhibit distinct behaviors (**Fig. 4H,I**). With ultra weak stickiness (bond strengths much lower than the thermal energy of the system), blue beads engaged in transient interactions that occasionally caused clumping (**Fig. 4J**), interspersed with frequent excursions to more dispersed states not easily distinguished from the free polymers (**Fig. 4I-J and Supp. Movies 1-2**). Quantifying the distribution of compaction and separation, we found broad distributions in the R_g_ (compaction) of the initially blue region compared to the initially gray region (**Fig. 4L**). The normalized distance of blue-to-blue vs. blue to gray (separation) were similarly broad **(Fig. 4L**), and both of these distributions overlapped the broad distributions seen with non-sticky blue beads (**Fig. 4L**). This organization is reminiscent of the behavior of the Pc-domains observed in the ORCA images, so for brevity we refer to this regime of the model as the Pc-chromatin-like regime.

Increasing the stickiness further produced a coil-to-globule phase transition (*37*), in which the blue region adopted a compact, largely spherical organization, that excluded gray portions of the polymer from the interior of this spheroid (**Fig. 4K**). This behavior is reminiscent of previous chromatin models based on homotypic polymer interactions (*15, 37, 48, 49, 55, 60–64*). Within the globule, 3D interactions between neighboring blue beads were transient, and beads rapidly exchanged 3D neighbors while maintaining a spheroidal organization – much like molecules in a liquid droplet (even though the beads are still chained together in a polymer, not a true liquid). For simplicity we refer to this behavior as the droplet regime for short. The stickiness in these simulations is still weak relative to the thermal energy of the system, (hence the transient nature of an individual bond) but collectively strong enough to maintain the spheroidal droplet throughout the simulation. In contrast to the previous “ultra-weak” interactions, the blue domain’s R_g_ was always smaller than that of the gray control region (**Fig. 4L** and **Supp Movie 3**). Similarly, the normalized blue-blue distances were always smaller than the normalized blue-gray distances (**Fig. 4L** and **Supp Movie 3**), contrasting the behavior in our data (**Fig. 3F**).

While clearly distinct at the level of single traces, at the population average level, the Pc-chromatin-like regime and droplet regime exhibited similar behaviors. Both formed a topologically associated domain (TAD) in the average contact-frequency plots, which aligned to the blue-region, though the absolute interaction frequency was higher in the droplet case and the domain extended slightly past the original boundaries (**Supp. Fig. 8**). Both also exhibited similar average bead-state profiles “in silico ChIP-seq” (**Fig. 4F** and **G**, low panels). By contrast, simulations with no sticky interactions exhibited no state-specific average 3D contact pattern across the population (**Supp. Fig. 8**), along with more uniform, lower average in silico ChIP-seq (**Fig. 4H,I**).

This averaging masks a few notable differences in how 3D structure relates to chromatin-state propagation in single cells. While maintaining a largely contiguous block of blue beads, the Pc-chromatin-like regime has a mild tendency for the whole block to drift upstream or downstream (**Fig. 4F**). In the droplet regime, we observed a tendency for flanking sequences, which erroneously acquired blue states, to rapidly fuse with the primary droplet and maintain their aberrant blue identity throughout the simulation (**Fig. 4G, red box**). As these fusions occur stochastically, their positional identity is largely averaged out across multiple independent simulations, though they do contribute to the mild expansion of the average TAD (**Supp. Fig. 8**). In contrast, in the Pc-chromatin-like regime, when blue states initiated ectopically, they were not stabilized and quickly disappeared (**Fig. 4F**). Thus, while both regimes are able to faithfully maintain the epigenetic memory, the Pc-chromatin-like regime is more robust at ejecting aberrantly marked domains.

The above simulations build a minimal model where 3D structural flexibility becomes important for maintaining fidelity of the Pc-repressive state in a dynamic nuclear environment. These simulations make several testable predictions about how changes in cellular properties should affect the 3D organization of the domain and the stability of repression (**Fig. 5A**). Cervical neural progenitor cells (cNPCs) have a Pc-target region (H3K27me3 domain) about half the length as that in brain cells, owing to the activation of the more anterior hox genes *a1-a5* (*18*) (**Fig. 5B**). Setting aside other differences in these cell types, the model predicts that the Pc-target region in cNPCs should be shifted to smaller R_g_-ratios and smaller normalized-distance ratios than observed in brain cells. To benchmark this shift, we first computed the compaction ratio and separation ratio from a non-self-interacting polymer and one in the droplet state. In the droplet case, these distributions lie completely to the right of 1, whereas for a non-interacting polymer they are both evenly distributed around 1 (**Fig. 5C**, note log-scale, **Supp. Fig. 9A,E**). Shortening the domain length by half reduced the fraction of the population with a compaction ratio greater than 1 from 81% to 57% (**Fig. 5D, Supp. Fig. 9C,F**). ORCA traces from cNPCs also shifted in the same direction, dropping from 82% to 61% (**Fig. 5E**). A similar agreement in the direction of the shift between simulations and ORCA measurements was observed in the fraction of cells with a separation ratio greater than 1 (**Fig. 5D-E, Supp. Fig. 9C, G**).

**Fig 5.**
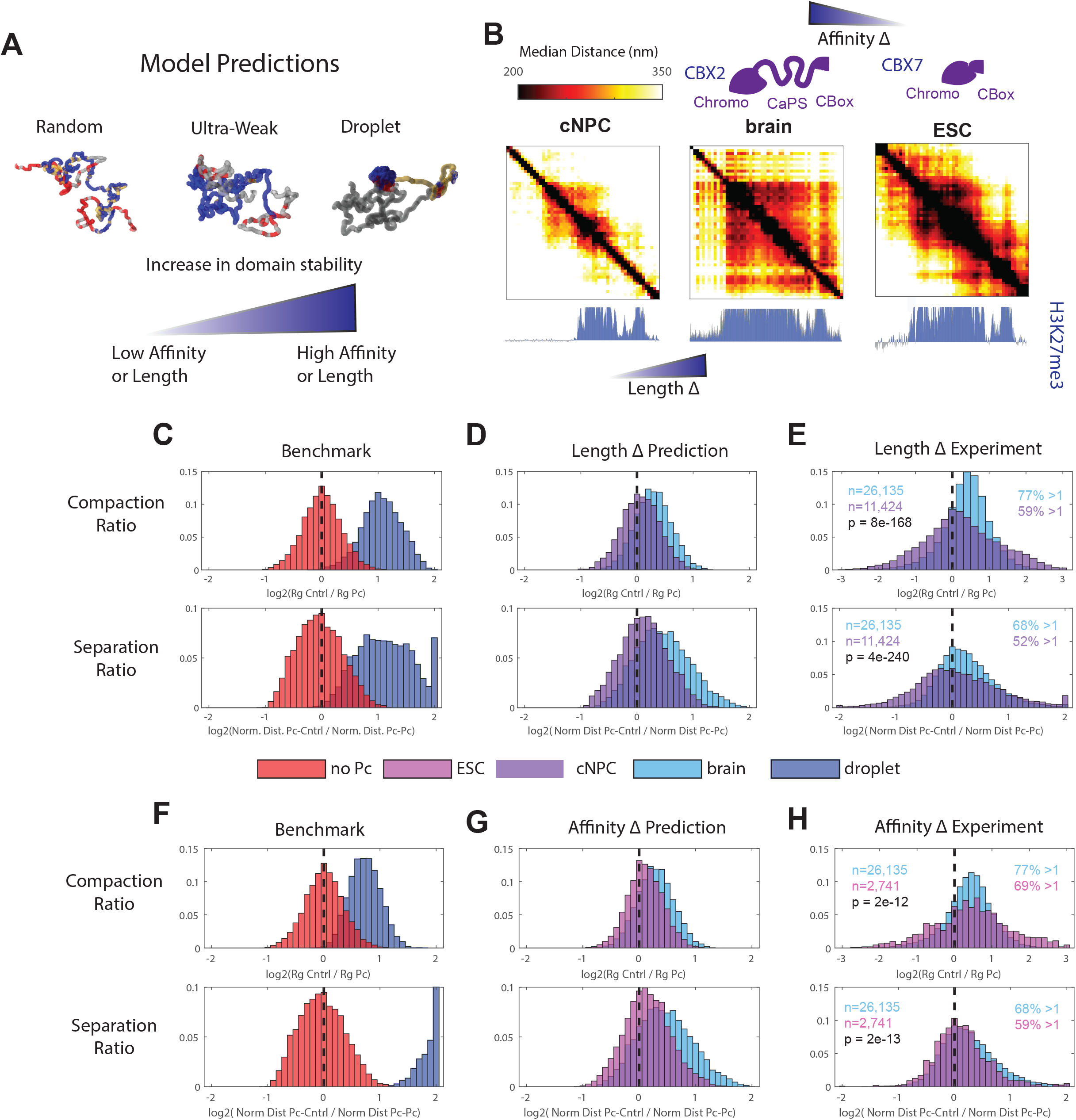
Length of a domain and the affinity of Polycomb subunits alters spatial feedback at repressive domains. **(A)** Schematic illustrating how modulating the affinity and domain length parameters of the model changes polymer structure. **(B)** Median distance plots of the experimental conditions for comparison with model predictions. Cervical neural progenitor cells (cNPCs) have smaller Pc-bound *Hoxa* domain (spanning only a6-a13) compared to brain cells. Mouse embryonic stem cells (mESCs) have a lower affinity CBX subunit compared to brain cells. **(C)** Top: histogram of the relative compaction (ratio of the Rg of the non-Pc domain to the Rg of the Pc-domain) in simulations of free polymers, and a droplet-forming PcG domain (as hypothesized in some prior models of Pc-domain folding). Bottom: separation ratio for the free-polymer and droplet simulation conditions. **(D)** Top: as in C but for simulations of polymers with a full length PcG domain (as found in brain cells), a half-length PcG domain (as found in cNPC cells). Bottom: as in C but for polymers with a full length PcG domain (as found in brain cells), a half-length PcG domain (as found in cNPC cells). (**E)** Top: Experimental data for the relative compaction in brain cells, in which the whole complex is Pc-repressed, vs. NPC cells, in which only half the complex is Pc-repressed. Bottom: The separation ratio for the same traces. **(F)** Simulation results plotted in C. **(G)** Simulation results comparing two full length domains with higher (brain) or lower (ESC) degrees of PcG-PcG affinity. **(H)** As in D, but contrasting compaction and separation for brain cells, which express the CBX2 PcG factor with high self-affinity, to ES cells, which express the CBX7 PcG factor instead, which has lower self-affinity.

The model also predicts a shift in compaction ratio and separation ratio for embryonic stem cells compared to brain cells. CBX2 is a core PcG factor with a large intrinsically disordered domain, which is essential for the ability of this protein to form liquid aggregates *in vitro* and *in vivo* and to compact nucleosome arrays *in vitro* (*8*). In embryonic stem cells, CBX2 expression is largely replaced by expression of a homolog, CBX7, which lacks this compaction and phase-separation (CaPS) domain (*8*). Thus, the model predicts that this removal of one of several “sticky” interactions among PcGs should shift the ratios in ESCs to lower values than exhibited by brain cells. We tested this by comparing the effect of changing stickiness at intermediate levels between those for the free polymer and droplet forming case (**Fig. 5F-G, Supp. Fig. 9A,E**). We observed in the ORCA data from ESCs, a shift to lower compaction ratios from 77% greater-than-1 down to 69%. This is the same direction we observed upon reducing the blue-bead affinity in our simulations (81% down to 63%) (**Fig. 5G-H**). This agreement in the direction of the shift is mirrored in the change in separation ratios (**Fig. 5G-H, Supp. Fig. 9B,D**).

Thus, the minimal model of Pc-chromatin structure and biochemistry provides both a possible explanation for its distinctive, yet non-compact structure, and provides a few experimentally testable predictions about structural change across cell types. The observation of varied degrees of spatial feedback between cell types examined experimentally suggests the possibility for cell-type specific diversification of domain stability. The stronger the effect of Pc-state on the 3D structure, the stronger the degree of feedback, and the more stable the epigenetic memory (**Supp. Fig. 10**).

## Discussion

By utilizing ORCA, combined with dynamic polymer simulations, we revisited the question of how PcG proteins organize chromatin and how this organization contributes to repression and epigenetic memory. We mapped the distribution of structural configurations across tens of thousands of single cells and explored how these distributions shifted between cell types and perturbations, observing substantial diversity and complexity of 3D configurations not predicted by prior models.

Prior models provided intuitive pictures of how Pc-chromatin structure (via compaction and/or separation) drives function (repression). The compaction model emphasizes the steric occlusion of bulky transcription machinery, drawing inspiration from the steric exclusion created by wrapping DNA on the nucleosome (*65*) (though the difference in scale between these two phenomena should not be overlooked). The high-frequency contact domains, seen in high-resolution Hi-C of Pc-domains (*18, 20, 66–68*); the compact structures seen in prior super-resolution microscopy (*15–19*); and the close clustering of short, in vitro nucleosomal arrays in the presence of several PcG proteins (*11, 35, 36*); all suggest entire Pc-domains (100s-1000s of nucleosomes) form heterochromatin that provides a steric barrier to transcription. While the canonical view of heterochromatin as a stable solid has increasingly given way to a more dynamic view, in which individual protein constituents rapidly turn-over and nucleosomes move within the compact region (*38*), the overall compact nature of the chromatin aggregate (amidst this molecular motion) is still viewed as a persistent state underlying the persistent silencing (*9, 38*). An alternative model emphasized the separation of Pc-chromatin from other chromatin, without insisting that the nucleosome arrangement is necessarily more dense or the mass-per-unit-volume any different. Separation through a liquid-liquid phase-separation (LLPS) process could segregate Pc-chromatin from active machinery, providing a chemical filter to exclude transcription machinery rather than a steric one (*7, 8, 38*). Such membraneless-organelle barriers, driven by LLPS, have now been described for a variety of subcellular processes (*69, 70*). This model drew support from the striking ability of PcG proteins to form liquid droplets that concentrate protein, permit rapid exchange of protein with solute outside the droplet, and allow droplet fusion (*39, 40, 71*), in a manner similar to nucleolar proteins (*72*). The decades old observation of the tendency of some PcG proteins to form visibly distinct foci in cells, called “Polycomb bodies”, are also suggestive of a membraneless-organelle (*6, 9*). The enhanced separation of Pc-domains from neighboring active chromatin, observed in super-resolution microscopy, also supports this separation model (*15, 17*). The two models are not exclusive, simple simulations based on weak multivalent interactions produce a regime which is both separated and compact (*15*). However, none of these models are consistent with the microscopy data we present here.

Many individual snapshots of Pc-repressed *Hoxa* domains we observed exhibited a degree of compaction and separation indistinguishable from that observed in non-Pc chromatin. Thus we reject the view that Pc heterochromatinizes its canonical targets, such as *Hoxa*, in order to silence them. Interactions that span only dinucleosomes (as probed in recent cryo-EM) (*73, 74*) or the short arrays of 4-12 nucleosomes (as probed in EM assays chromatin compaction (*11, 35, 36*)Should these clutch size, Pc-dependent multi-nucleosome interactions exist in vivo chromatin (as we believe likely), we propose they reflect transient interactions. The biological role of these transient interactions may have more to do with adding a degree of chromatin 3D spatial-feedback to the chemical feedback, rather than enabling a steric mechanism of repression akin to that achieved by DNA-nucleosome interactions. Recent structural analysis supports this view, which emphasizes the potential for catalytic stimulation between juxtaposed PRC2 complexes (*73*). As such, we propose it is better not to describe the organization of Pc-chromatin as “compact” or as “heterochromatin”, which invoke steric roles for the structure that are difficult to reconcile with single molecule data.

The Pc-chromatin domains we observed also lacked the sort of spatial separation from neighboring chromatin expected for chromatin segregated into a membraneless organelle, as happens with rDNA segregation into the nucleolus. Individual portions of the Pc-domain were not infrequently closer to non-Pc chromatin than to Pc-chromatin, and droplet structures were exceedingly rare, and probably transient in the data. This is in stark contrast to the type of 3D organization we observe in simulation of chromatin droplet formation. This does not rule out an essential role in Pc-repression for the IDRs and multerimerization domains of PcG proteins, features which also play critical roles in their ability to form liquid-protein droplets. Indeed, in our modeling efforts to explain the complex, heterogeneous structures, we observe a key roles for such interactions. However, rather than a narrow view of the emergent properties of weak, multivalent interactions among disordered protein structures being limited to their ability to produce first order phase transitions (intermixed to separated) - our data indicate the complex interactions among PcG proteins at target chromatin operate in a regime that allows a more complex array of 3D structural ensembles (differing between Pc-degraded ESCs, ESCs, NPCs, and brain). This gradation of compaction and separation is not representative of condensate formation, where-in a critical concentration of factors is reached causing an abrupt collapse into a compact, largely spherical globule (*37*).

This suggests development may tune the nature of Pc-repression to balance stability of silencing against potential for reprogramming. The larger the effect of the Pc state on 3D structure (compared to the Pc-free polymer), the stronger the spatial feedback. Cells with minimal spatial feedback (e.g. low sticky model / mESCs data) will be more susceptible to breakdown of the repressed domain. This reduced stability against random perturbation also implies a reduced stability against developmentally queued perturbations, making these cells more amenable to converting part or all of the domain out of the Pc-repressed state. Meanwhile, cells with a higher degree of spatial feedback (e.g. high sticky model / brain cell data) will maintain more stable repression, but be harder to reprogram. Consistent with this, mice carrying a mutant PcG, CBX2, that lack a self-sticky, disordered region of the protein, exhibit partially penetrate anterior-posterior homeotic transformations of the skeleton (*36*), indicative of stochastic loss of Pc-repressive state in some cells. Similarly, introducing CBX2 expression in embryonic stem cells, in place of the less self-sticky CBX7 homolog that is normally expressed, impaired differentiation (*75*). Other cell-type specific and locus-specific variation in PcG composition that alter the effective strength of PcG-PcG interactions may similarly tune between stability and reprogramming in a cell-type or locus specific manner, while still providing a degree of repression and accessibility less expected in a liquid condensate forming regime.

While we find the data incompatible with a model in which the unique 3D structure of Pc-chromatin creates a steric (compact) or a chemical-surface (liquid droplet) barrier to transcriptional machinery, our simulations demonstrate one alternative way the 3D structure could be important to Pc-repression, that is consistent with our data. This third model is that by shaping the 3D structure of the repressed chromatin, Pc provides spatial feedback to couple with the known biochemical feedback and stabilize the epigenetic memory. By incorporating the multi-state nature of Pc-chromatin into our simulations we find a regime in which the structures may make frequent excursions into decompact and intermixed states, (as also seen in the distribution of structures in our ORCA data) and still provide a major boost to the domain stability compared to a system lacking spatial feedback. The simulations also identified a liability of the droplet-forming regime examined in prior models (*44, 48, 49*), which is at greater risk of stabilizing ectopic domains. In our model, spatial and biochemical feedback combine to ensure robust epigenetic inheritance. Repression of transcription is assumed to be a property of protein occupancy (state) rather than something achieved directly through the higher-order (kilobase-scale) structure. This view is consistent with the behavior of other transcription factor repressors, which achieve efficient silencing without detected changes to high-order chromatin folding.

The complex distribution of structures adopted by Pc-target chromatin in our data, were largely reproduced by a minimal model that did not require explicit representation of many known features of Pc-repression in general and at *Hox* loci in particular. It was unexpected and striking that the model was able to retain stable boundaries without requiring biochemical and genetic elements previously associated with epigenetic boundaries. Both genetic and biochemical analyses have uncovered prominent roles for Trithorax group (TxG) proteins in PcG regulation, which oppose the activity and spreading PcG proteins and generally facilitate gene activation rather than repression (*29, 76*). Prior modeling work has already shown how competition between self-reinforcing, mutually antagonistic systems, such as PcG/TxG competition, can create stable boundaries without reference to 3D chromatin structure (*50, 52*) (provided turnover is limited and non-specific recruitment non-existent). We expect such competition would reinforce the stability of PcG domains, but find it interesting that relatively stable boundaries can arise without it. Similarly, we find it is not necessary to assume the existence of sequence specific bias in PcG recruitment within the domain, though such biases are known to play a role in recruitment and memory in flies and mammals (*1, 5, 77*). We can speculate that such sequence-specific elements may strengthen boundaries, in some cases through a TxG dependent mechanism, given their roles in recruiting both factors (*29*). The non-requirement for sequence-specific elements in domain stability, as seen in our modeling, is consistent with recent observations that Polycomb Response Elements from the engrailed complex - some of the most studied sequence-specific PcG targeting elements known, may be mutated at their endogenous target without disrupting stable PcG-domain maintenance (*78*). At borders of repressed chromatin, and *Hox* loci in particular, the importance of CTCF in maintaining stable borders and preventing loss of PcG-chromatin is established in both flies (*79*) and mammals (*20, 67*). Sequence specific borders provide an intuitive way to stop nearest-neighbor spreading, and likely can reinforce the intrinsic border memory created by the 3D structure of PcG domains - but are not required. We note that in the pluripotent ES state or the differentiated embryonic brain, the *Hoxa* Pc-domain downstream (left) boundary is not delineated by either CTCF or TxG marks. Interestingly, the smaller Polycomb domain observed in the NP state, which the model predicts should be less stable than in the brain, is delineated by both CTCF and TxG marks.

From these data and simulations, we propose the distinctive 3D structure of Polycomb-domains functions as a spatial feedback system. This spatial feedback is coupled to known biochemical feedback, and enables portions of the sequence to reinforce Pc-chromatin state at other portions of the sequence which start to lose it. This process requires flexibility of the polymer, not a regular structure as found in many folded proteins. Nothing in this mechanism of spatial feedback is incompatible with maintenance of an epigenetic state of activation instead of repression, despite the fact it contributes to a statistical tendency of the domain to be “compact” (smaller in size), a chromatin feature often attributed a repressive role. While simulations show that larger domains may undergo a more binary transition from a random-walk polymer to a membraneless nuclear organelle, the data suggest that Hox loci involve too few interactions, and exhibit instead a more gradual transition between these states. We speculate that this graded transition provides an opportunity for evolution to strike a balance between the stability of repression and the ability to reprogram targeted portions of the domain into an active state in a cell-type dependent manner.

## Acknowledgements

We thank Joe Wakim and Andrew Spakowitz for detailed discussions and feedback on modeling chromatin polymers and PcG interactions. We thank TzuChiao Hung and Mervenaz Koska for generously providing the mouse brain and embryo tissues imaged in this study. We thank Andrew Fire, Anne Villeneuve, Andy Spakowitz and Lacra Bintu for critical feedback during these investigations and a critical reading of the manuscript. The work was supported by a Stanford Bio-X Fellowship to S.E.M., and grant EF2022182 from the N.S.F, grant U01DK127419 from the N.I.H., a Beckman Young Investigator Award and Packard Fellowship to A.N.B.

## Supplemental Figure Legends

**Supp. Fig. 1.**
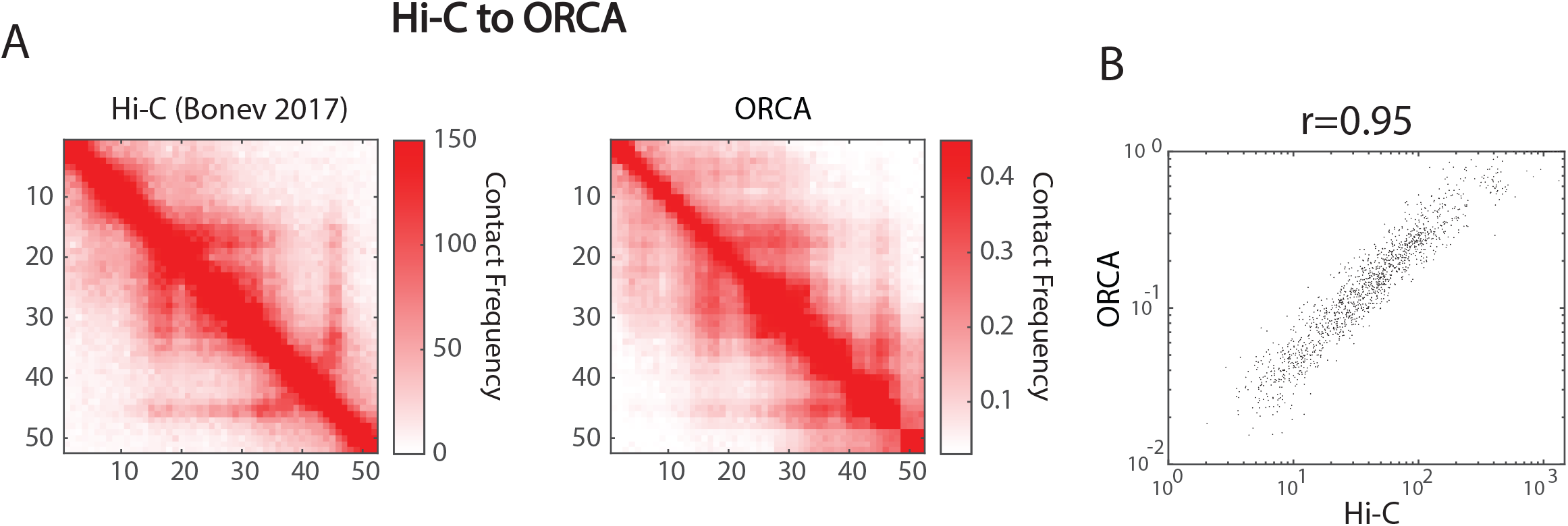
Comparison of Hi-C and ORCA. **A** Hi-C map at 5kb resolution of the mES Hox-A locus is qualitatively similar to the contact frequency measured with ORCA (contact defined by a 150 nm threshold). **B** The two techniques produce correlated interaction frequencies (r=0.95).

**Supp. Fig 2.**
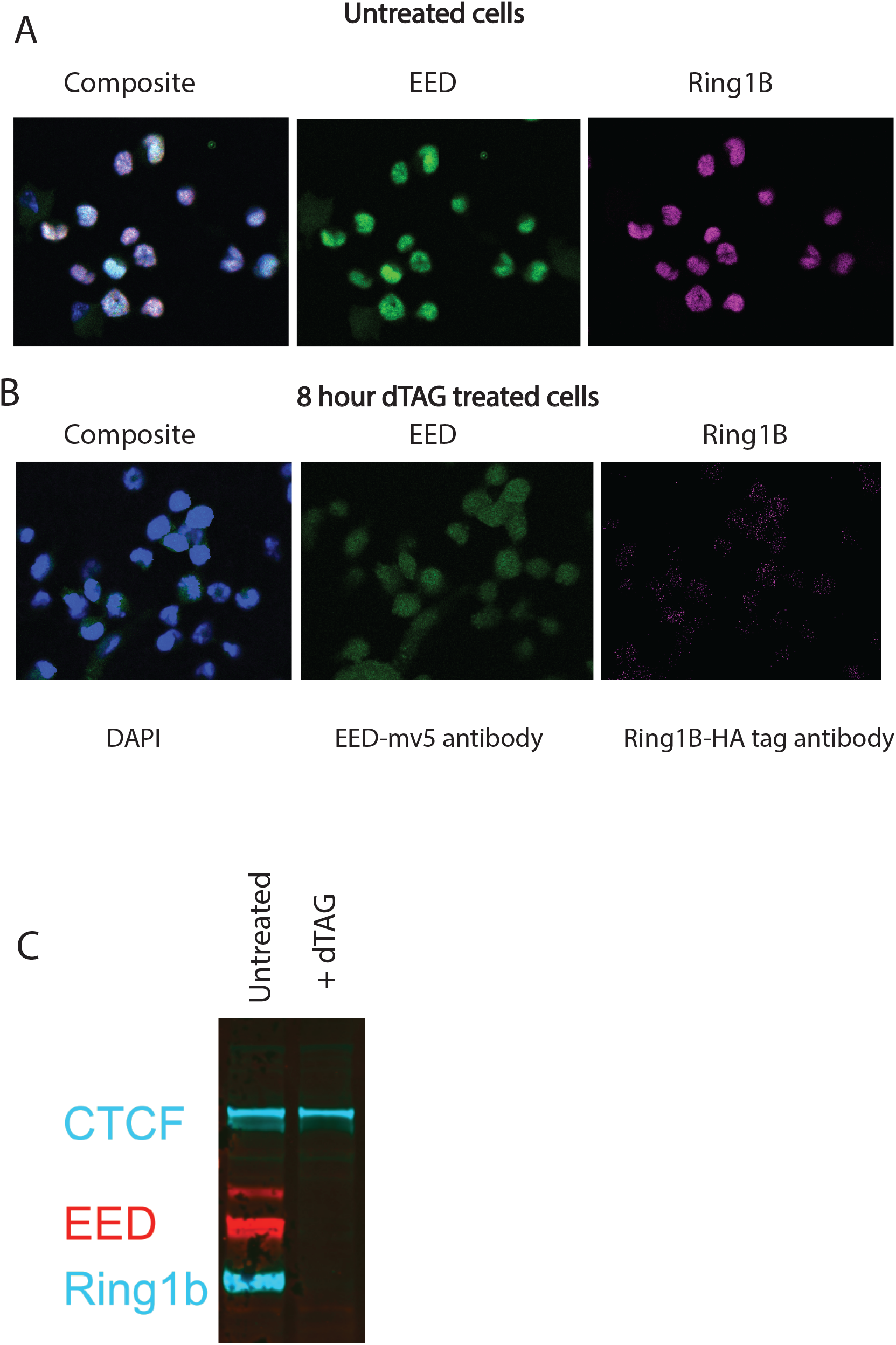
Degradation of Ring1b and EED targets with dTAG ligand. **A** Immunofluorescence of untreated cells labeled with antibodies against v5 tag (Ring1b-v5 tagged) and HA tag (EED-HA tagged). **B** Immunofluorescence of cells after 8 hours of treatment with dTAG ligand labeled with antibodies against v5 tag (Ring1b-v5 tagged) and HA tag (EED-HA tagged). **C** Western blot of untreated vs. 8 hours of dTAG treatment.

**Supp. Fig 3.**
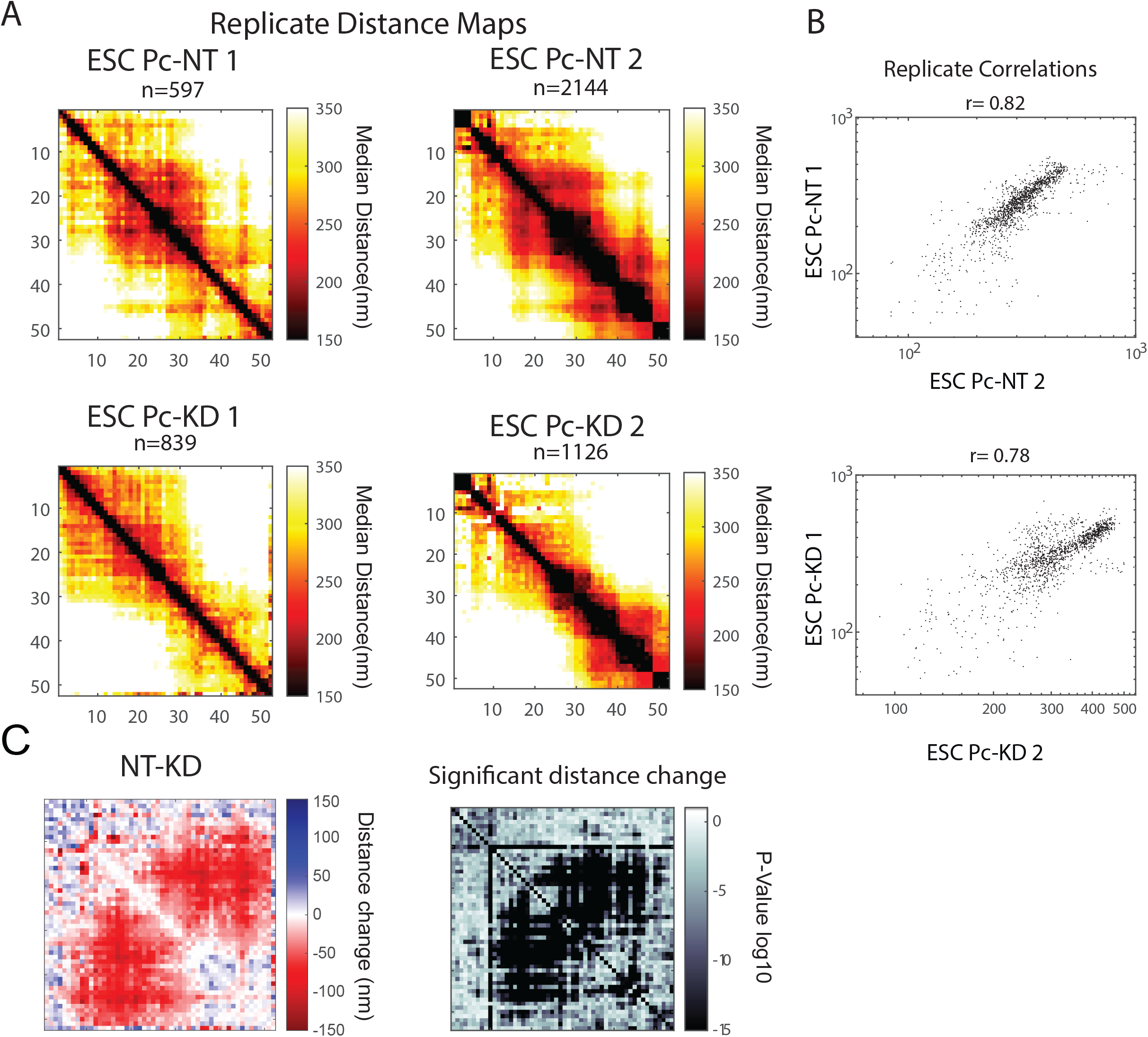
Reproducibility of mESC results and PcG depletion results. **A** Median distance plots of each replicate of the no treatment mES cells and dTAG treated mES cells **B** Correlation plots of median distances between the no treatment mES cells and dTAG treatment mES cell replicates **C** Subtraction plot of the no treatment - dTAG treated replicates, significant distance change (p << 0.05) shown in plot where darker area signifies greater significance

**Supp. Fig. 4.**
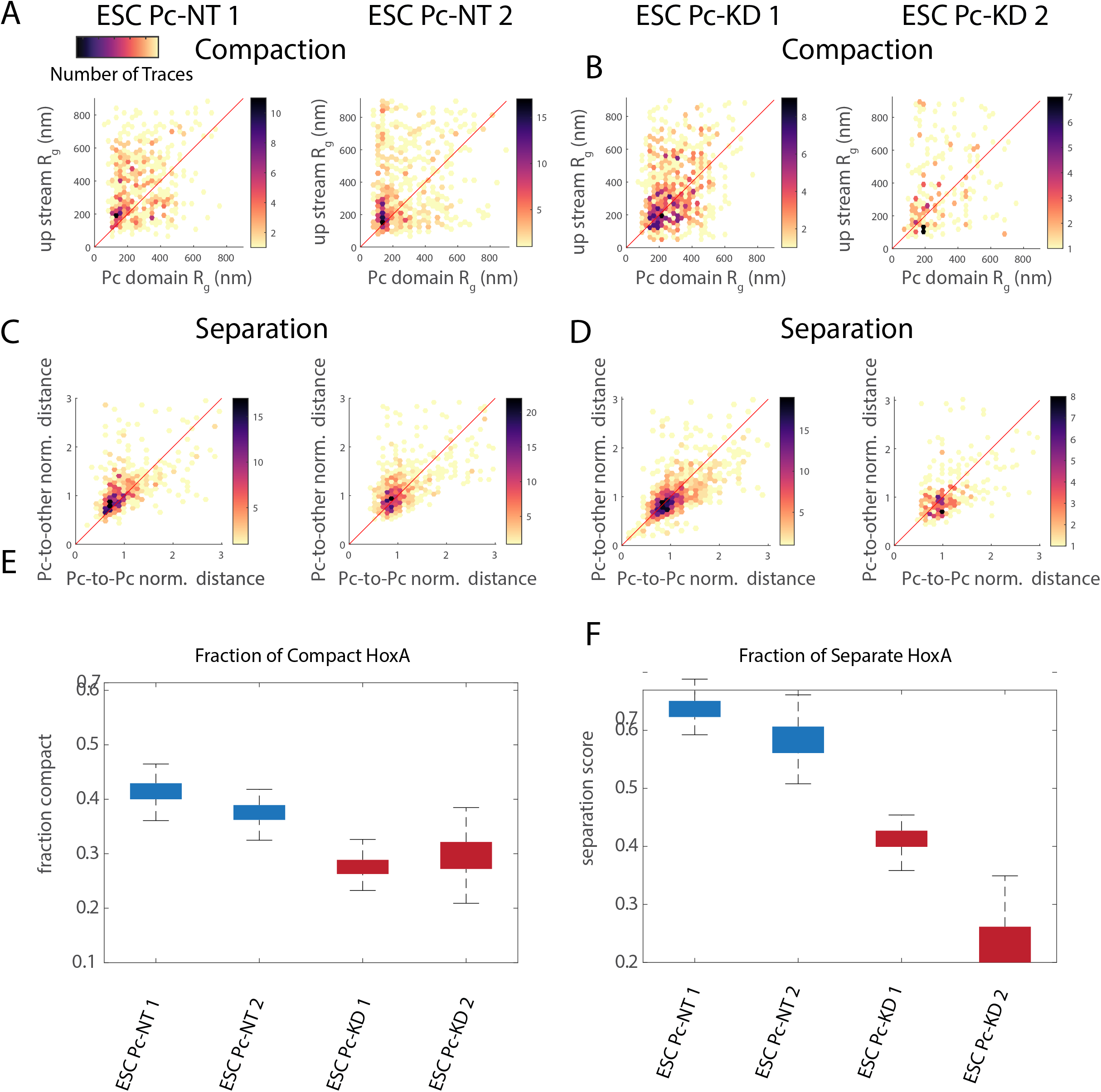
Replicate comparison of compaction and separation results. **A** Replicate 2D compaction plots for untreated mES cells. **B** Replicate 2D compaction plots for the dTAG treated mES cells. **C** Replicate 2D separation plots for untreated mES cells. **D** Replicate 2D separation plots for dTAG treated mES cell. **E** Summary box plots for fraction of compact Hoxa loci

**Supp. Fig 5.**
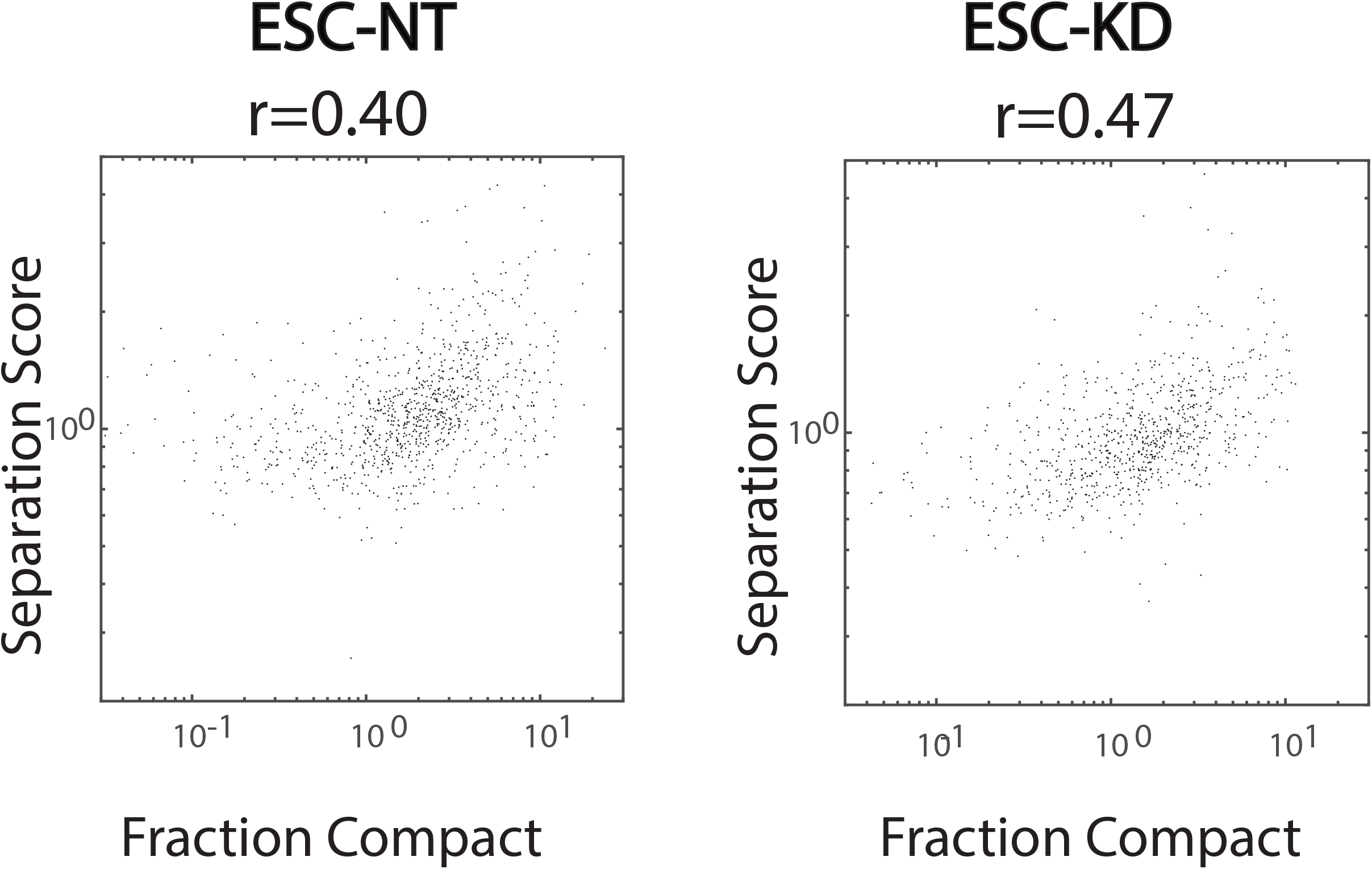
Correlation between separation and compaction measurements. Correlation between compaction and separation measurements in untreated and knockdown samples.

**Supp. Fig. 6.**
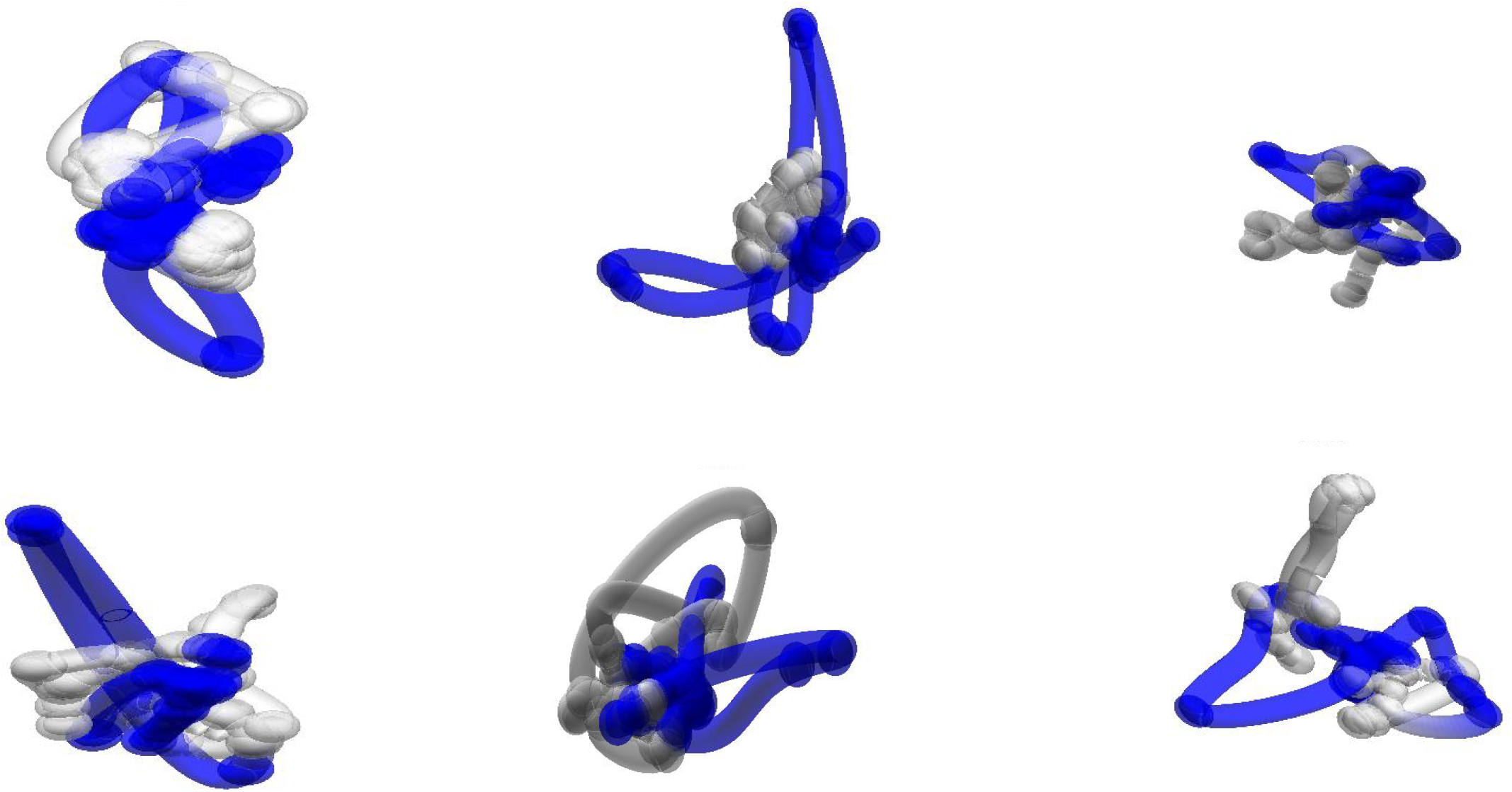
polycomb knockdown example traces. Example traces from cells following PcG depletion

**Supp Fig. 7.**
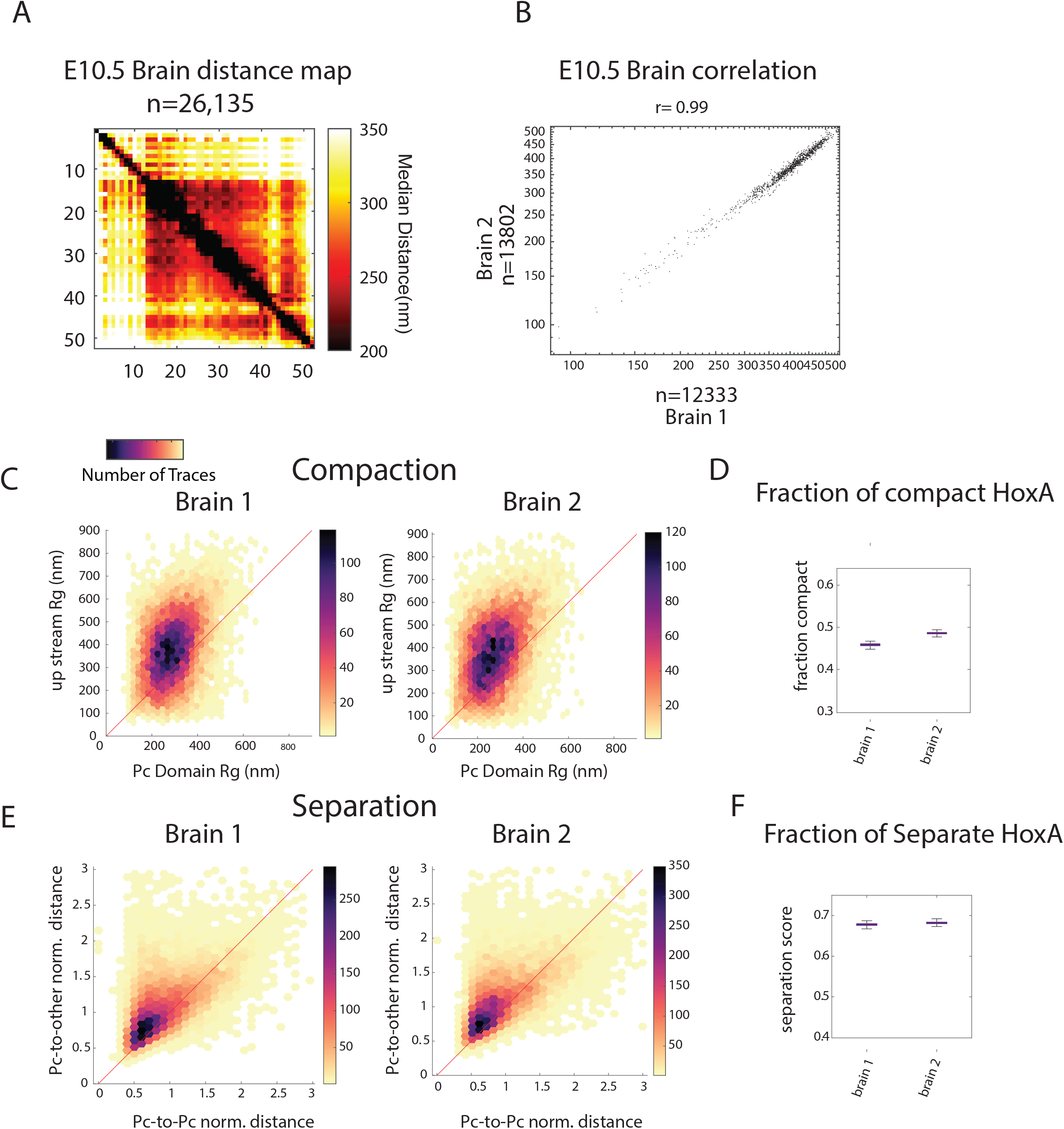
Reproducibility of data between embryonic brains. **A** Median distance map of two E10.5 brain sections for the Hoxa domain. **B** Correlation plot of median distances for brain replicates (r=0.99). **C** 2D-plots of upstream Rg (nm) Vs. Pc domain Rg (nm) for each brain replicate. **D** Fraction of loci with compact Pc Domains for each brain replicate. **E** 2D-plots of normalized interdomain distance vs normalized Pc Domain distance for each brain replicate. **F** Fraction of loci with demixed Pc Domains for each brain replicate.

**Supp. Fig. 8.**
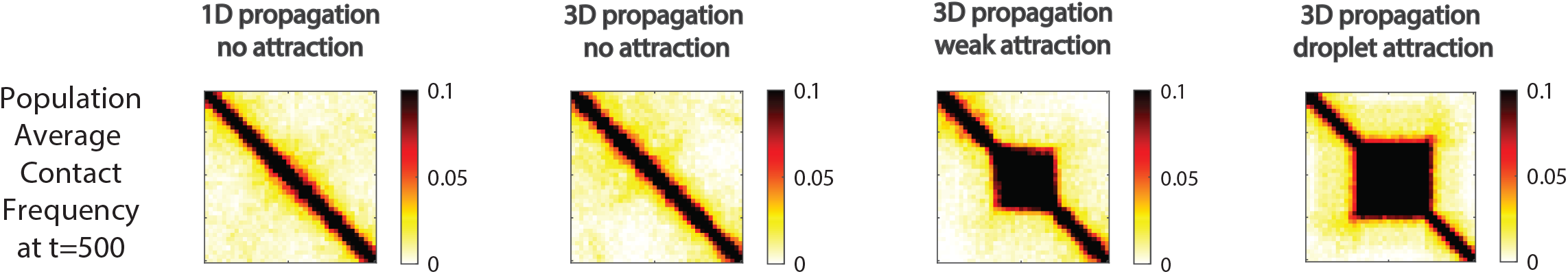
Contact Frequency. Population average contact frequency of each modeled condition.

**Supp. Fig. 9.**
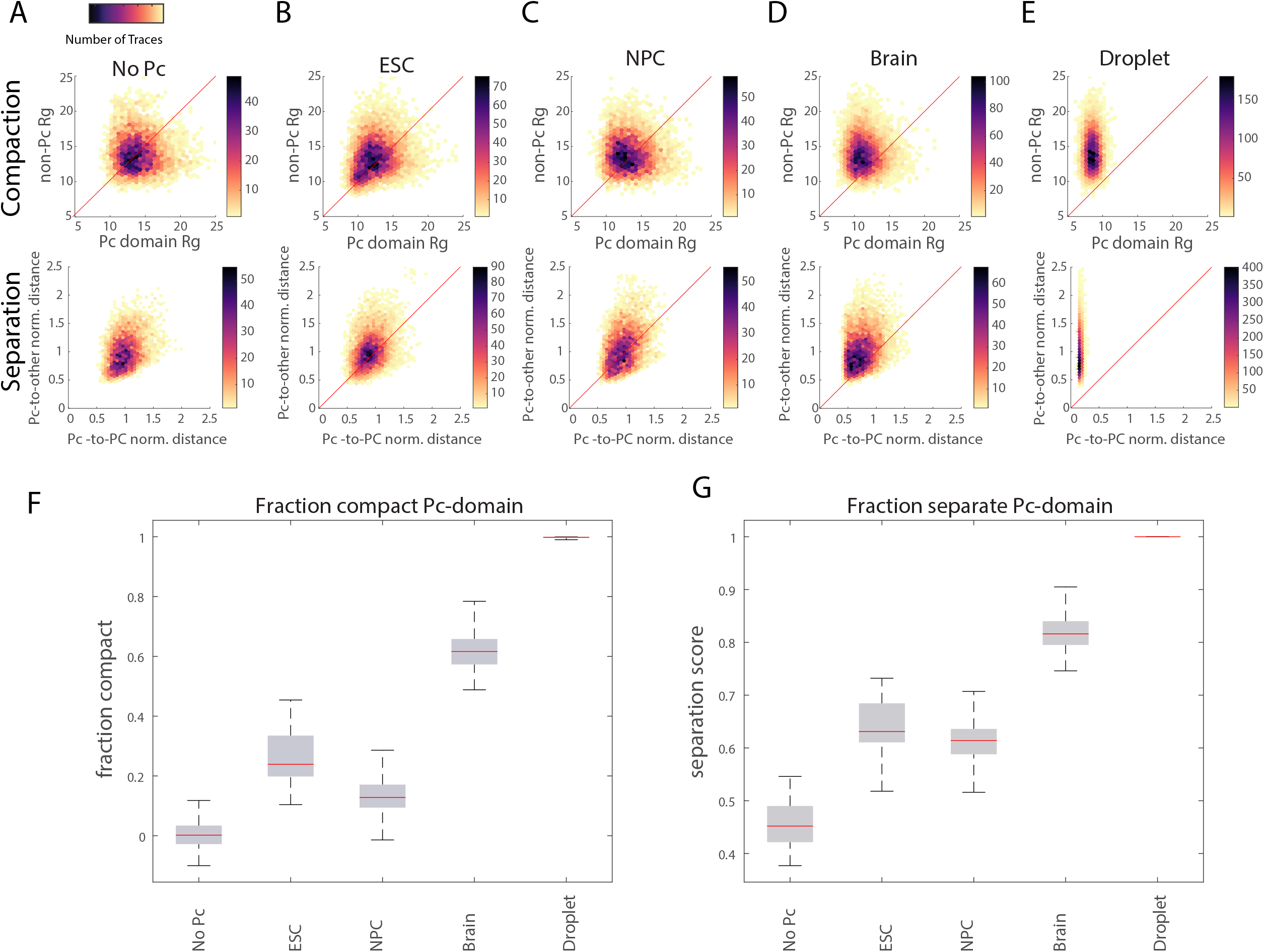
Compaction and separation analysis for simulated polymers. **A** 2D-compaction and separation plots for no-Pc/random polymer simulations with attractive strength of 0 and domain length of 200 monomers. **B** As in A but for interaction strengths of 0.3 to represent ESCs. **C** As in B but for interaction strength of 0.4 and a domain length of 100 monomers to simulate NPCs. **D** As in D but with domain length of 200 monomers. **E** As in D but with attraction strength of 0.75 simulating the droplet model. **F** Fraction of compact Pc-domains for the different simulations. **G** Fraction of separate Pc-domains for the various simulations. All simulations include added Gaussian random noise of variation 5 units to simulate measurement error, calibrated from the data using replicate labeling (∼50 nm).

**Supp. Fig. 10.**
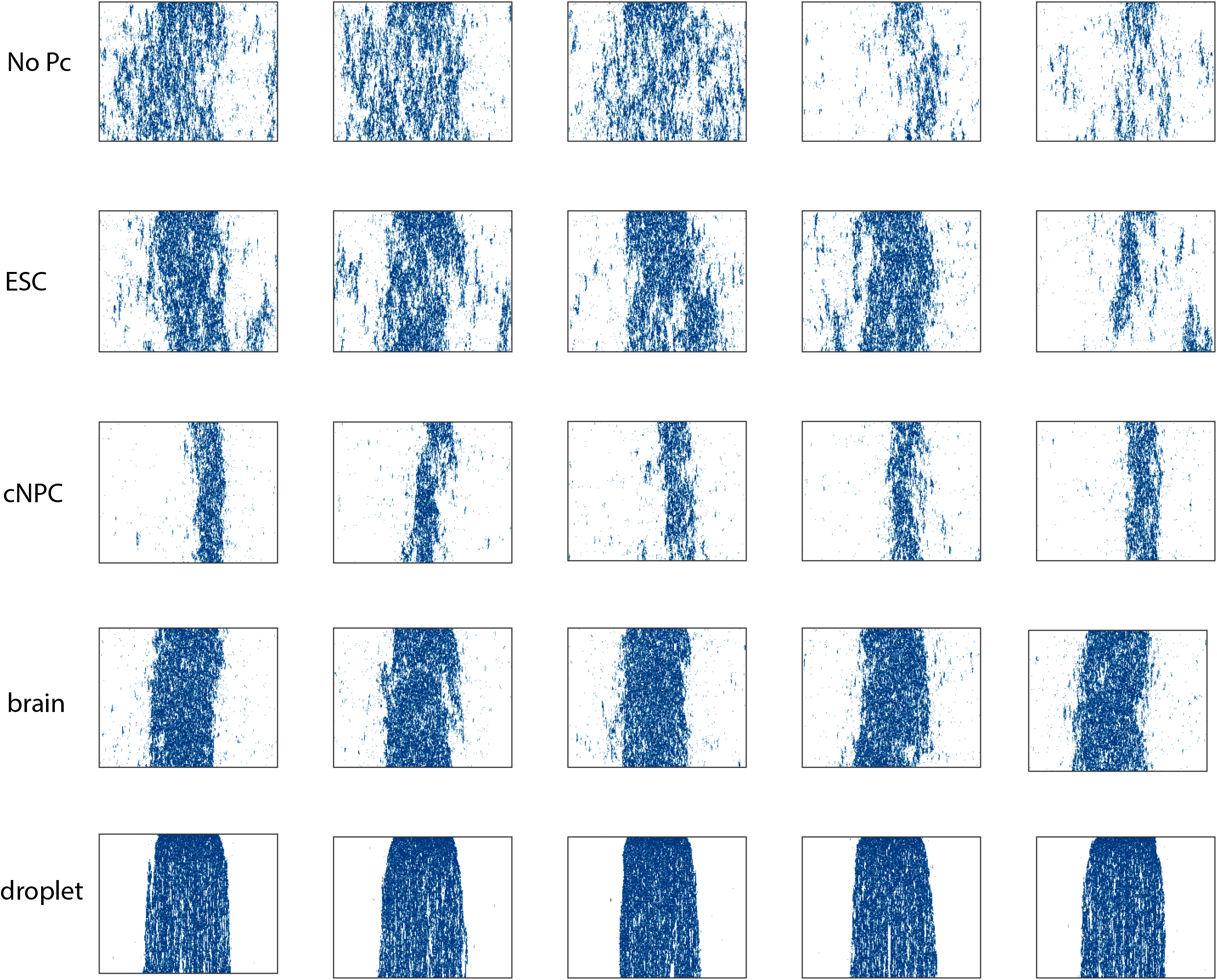
Domain stability increases with increased spatial feedback. Panels show kymographs, as in Fig 4, for examples from several independent simulations under each different conditions shown in Fig 5: free polymer (no Pc), simulated ESCs (lowest stickiness), simulated brain (higher stickiness), simulated cNPCs (shorter domains) simulated droplets.

## Supplemental movies

Supp. Movie 1: Simulation No Attraction:

[SimNoAttr_BlueRedYellow_3D1D.mp4](https://drive.google.com/file/d/17a7d3Ubl8U6AWsWAn36LS8aLuGySgN6L/view?usp=sharing)

Supp Movie 2: Simulation Dynamic Pc-domain structures exhibit mixed compaction and memory

[SimDynPc_BlueRedYellow_3D1D.mp4](https://drive.google.com/file/d/1VdDFEAuDtttpSTdaLkx14Qtx2B-G_ZxV/view?usp=sharing)

Supp Movie 3: Simulation Droplet-like Pc-domains stabilize ectopic Pc deposition

[SimDroplet_BlueRedYellow_3D1D.mp4](https://drive.google.com/file/d/1jMFH-TEHLWZUMKp7aqIBY3669vwwSHGU/view?usp=sharing)

## Notes

### Competing Interest Statement

The authors have declared no competing interest.

